# Recruitment and abundance of intertidal barnacles and mussels along the Atlantic Canadian coast: pelagic influences and relationships with predator abundance

**DOI:** 10.1101/239756

**Authors:** Ricardo A. Scrosati, Julius A. Ellrich

## Abstract

Benthic species from rocky intertidal systems are irregularly distributed along marine coastlines. Nearshore pelagic conditions often help to explain such variation, but most such studies have been done on eastern ocean boundary coasts. Through a large-scale mensurative study, we investigated possible benthic-pelagic coupling along the Atlantic coast of Nova Scotia, a western ocean boundary coast. We studied the high intertidal zone of nine wave-exposed bedrock locations spanning 415 km of coastline from north to south. At each location in the spring, we measured the recruitment of barnacles and mussels, the two main filter-feeding invertebrates. Recruitment varied irregularly along the coast. Satellite data on coastal phytoplankton and particulate organic carbon (food for intertidal filter-feeders and their pelagic larvae) and in-situ data on seawater temperature explained, to varying degrees, the geographic structure of recruitment. In turn, the summer abundance of both filter-feeders was positively related to their spring recruitment. Ultimately, predator (dogwhelk) abundance was positively related to the recruitment and/or abundance of barnacles and mussels (the main prey of dogwhelks). These results are consistent with bottom-up forcing influencing intertidal community structure on this coast. Sea ice may also influence this predator–prey interaction. Drift ice leaving the Gulf of St. Lawrence in late winter disturbed the northern locations surveyed on the Atlantic coast, making barnacles (owing to their high spring recruitment) the only food source for dogwhelks at such places. Investigating the oceanographic drivers of pelagic food supply and seawater temperature should help to further understand how this large metacommunity is organized.

## Introduction

Understanding the spatial variation of ecological systems is a central goal of ecology. Marine rocky shores have often been studied in this regard because of the relative ease of sampling to examine them. Concepts such as benthic-pelagic coupling and bottom-up forcing have commonly been advanced through research on rocky intertidal systems, for instance. Benthic-pelagic coupling refers to the influence of nearshore pelagic conditions on benthic coastal processes (Navarrete et al. 2005, Griffiths et al. 2017), while bottom-up forcing refers to the effects of food or nutrient supply on basal trophic levels that propagate through consumption to higher trophic levels (Menge 1992).

Pelagic conditions that affect intertidal systems change at various spatial scales. For example, at local scales, wave exposure differences due to topographic variation affect pelagic food supply (Steffania and Branch 2003, McQuaid and Lindsay 2007), larval dispersal (Bertness et al. 1992), and benthic survival (McQuaid and Lindsay 2000, Larsson and Jonsson 2006, D’Amours and Scheibling 2007), thereby influencing species distribution and abundance (Heaven and Scrosati 2008). At regional scales, nearshore pelagic traits such as seawater temperature and planktonic food abundance change for oceanographic reasons and generate regional patterns in intertidal species distribution and abundance (Menge et al. 2003, Blanchette et al. 2008, Shanks et al. 2017a). This paper is mainly concerned with advancing knowledge at regional scales.

Benthic-pelagic coupling and bottom-up forcing in rocky intertidal systems have often been studied by surveying wave-exposed habitats. Such habitats face the open ocean directly, which facilitates the identification of nearshore pelagic influences. Those studies have mainly been done on temperate shores, both in the northern (Menge et al. 2009) and southern (Navarrete et al. 2005) hemispheres. However, they have overwhelmingly been conducted on coasts from eastern ocean boundaries (Menge and Menge 2013). Such coasts frequently experience upwelling and are biologically productive, supporting important fisheries (Food and Agriculture Organization 2016). Those studies have concluded that oceanography-mediated changes in phytoplankton abundance and particulate organic carbon (food for intertidal filter-feeders) and sea surface temperature influence intertidal ecological patterns (Menge et al. 1997a, 2009, Wieters 2005, Shanks et al. 2017a). Influences can be direct (e.g., phytoplankton increasing the abundance of intertidal filter-feeders; Menge et al. 1997b) or indirect (e.g., phytoplankton increasing the abundance of intertidal predators through direct effects on their filter-feeding prey; Leonard et al. 1998). Although it is reasonable to expect similar relationships on other coasts, their magnitude has commonly not been quantified. This is the case for most coasts from western ocean boundaries. Addressing this gap is necessary to produce a more general predictive framework.

Studies of this kind have employed various approaches, including large-scale mensurative studies, local-scale experiments, or combinations of both (Menge and Menge 2013). For coasts on which limited knowledge exists, mensurative approaches are especially useful to identify basic regional patterns, a necessary precondition for more detailed studies (Underwood et al. 2000, Sagarin and Pauchard 2010). Using large-scale mensurative data, this paper provides for the first time evidence in support of benthic-pelagic coupling and bottom-up forcing along the Atlantic coast of Nova Scotia, Canada. There is a rich history of rocky intertidal research for this coast (McCook and Chapman 1997, Hunt and Scheibling 1998, Scrosati and Heaven 2007, and references therein), but studies have largely been restricted to a few locations near research institutions. For instance, there is no comprehensive account of latitudinal changes in intertidal invertebrate recruitment and abundance and the relationships with relevant pelagic traits (e.g., seawater temperature and food supply) and intertidal predator abundance.

As on other temperate shores across the world (Bustamante and Branch 1996, Navarrete and Castilla 2003, Blanchette and Gaines 2007, Nakaoka et al. 2006, Hawkins et al. 2009, Lathlean et al. 2010, Arribas et al. 2013, Menge and Menge 2013), barnacles and mussels are often dominant organisms in wave-exposed, rocky intertidal habitats in Nova Scotia (Hunt and Scheibling 1998, Scrosati and Heaven 2007). Because of the ecological importance of these filter-feeders, their recruitment is commonly measured to investigate benthic-pelagic links (Menge and Menge 2013, Mazzuco et al. 2015, Shanks and Morgan 2017). Therefore, the first objective of our study was to investigate the latitudinal variation in intertidal barnacle and mussel recruitment along the Atlantic coast of Nova Scotia. To seek evidence of benthic-pelagic coupling, we examined how barnacle and mussel recruitment was related to coastal seawater temperature, phytoplankton abundance, and particulate organic carbon. We then evaluated how barnacle and mussel recruitment was related to the abundance of these organisms later in the year. Finally, in search of signs of bottom-up forcing, we assessed whether the recruitment and abundance of barnacles and mussels were related to the abundance of their main predators (dogwhelks) along the coast.

## Materials and Methods

### Locations

For this study, we surveyed nine intertidal locations that span the full Atlantic coast of mainland Nova Scotia, nearly 415 km between Glasgow Head and Baccaro Point (Fig. 1). For ease of interpretation, the locations are referred to in the text as L1 to L9, from north to south. Their names and coordinates are provided in Table 1. These are wave-exposed locations, as they directly face the open waters of the Atlantic Ocean (Fig. 2). Values of daily maximum water velocity measured with dynamometers (see design in Bell and Denny 1994) in such habitats range between 6–12 m/s (Hunt and Scheibling 2001, Scrosati and Heaven 2007, Ellrich and Scrosati 2017). The studied locations consist all of stable bedrock.

**Figure 1.**
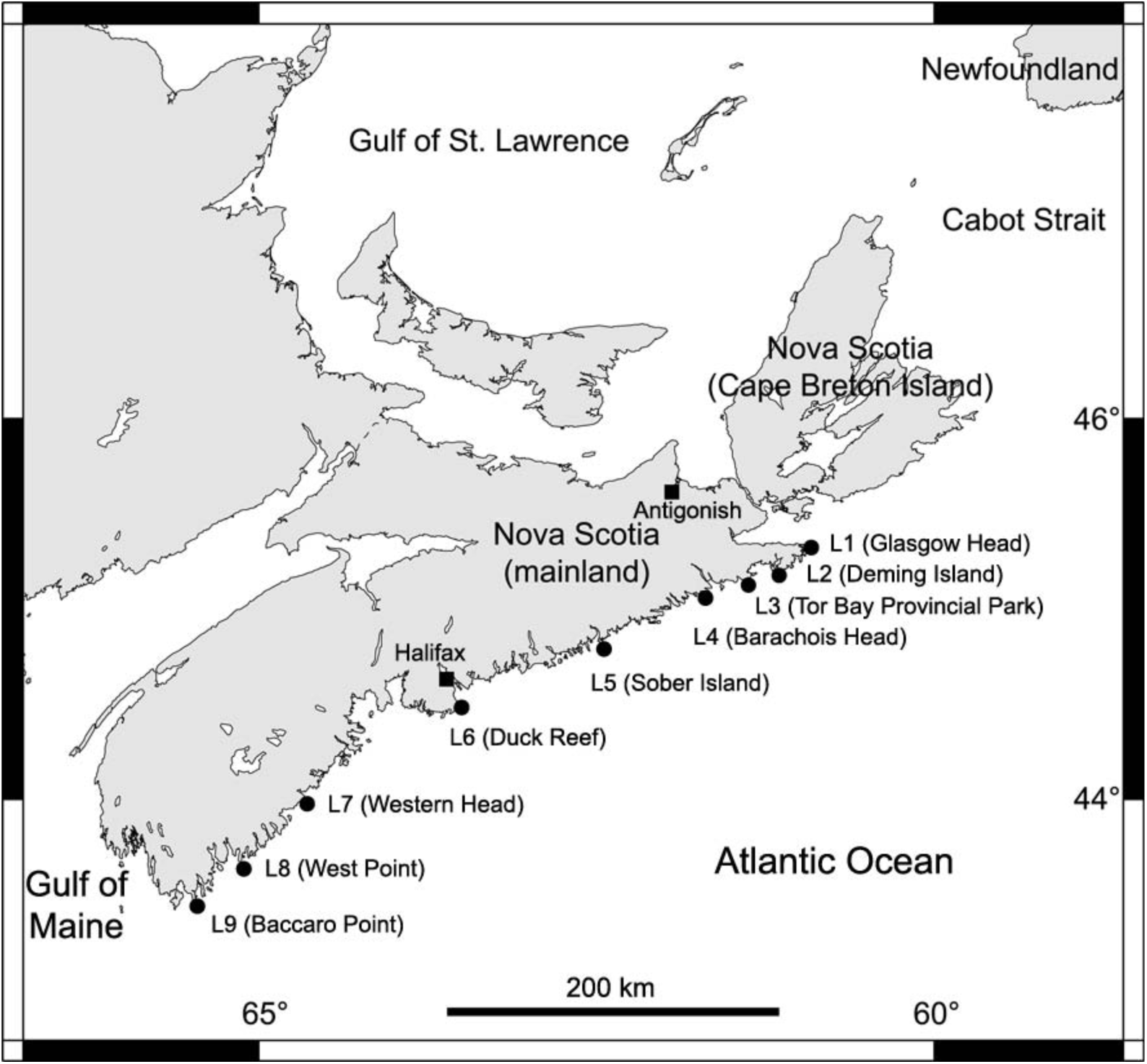
Map indicating the nine locations surveyed along the Atlantic coast of mainland Nova Scotia, Canada.

**Table 1.** Basic information about the nine wave-exposed locations surveyed for this study.

| Location code | Location name<br>(geographic coordinates) | Closest location with tide data (geographic coordinates) |
| --- | --- | --- |
| L1 | Glasgow Head<br>(45.3203, -60.9592) | Canso<br>(45.3500, -61.0000) |
| L2 | Deming Island<br>(45.2121, -61.1738) | Whitehead<br>(45.2333, -61.1833) |
| L3 | Tor Bay Provincial Park<br>(45.1823, -61.3553) | Larry's River<br>(45.2167, -61.3833) |
| L4 | Barachois Head<br>(45.0890, -61.6933) | Port Bickerton<br>(45.1000, -61.7333) |
| L5 | Sober Island<br>(44.8223, -62.4573) | Port Bickerton<br>(45.1000, -61.7333) |
| L6 | Duck Reef<br>(44.4913, -63.5270) | Sambro<br>(44.4833, -63.6000) |
| L7 | Western Head<br>(43.9896, -64.6607) | Liverpool<br>(44.0500, -64.7167) |
| L8 | West Point<br>(43.6533, -65.1309) | Lockport<br>(43.7000, -65.1167) |
| L9 | Baccaro Point<br>(43.4496, -65.4697) | Ingomar<br>(43.5667, -65.3333) |

**Figure 2.**
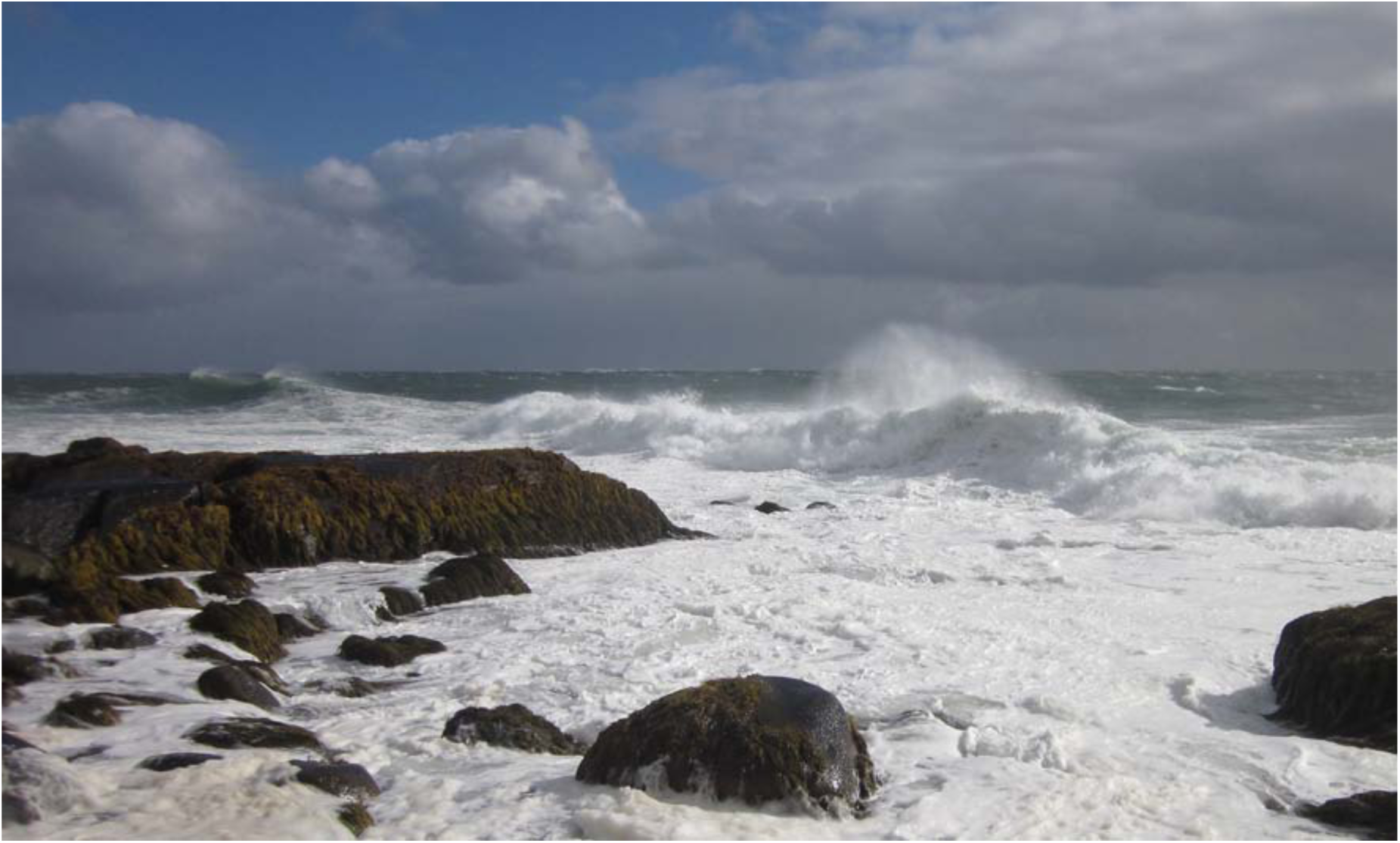
Typical view of wave-exposed habitats on the Atlantic coast of Nova Scotia. Photo taken by R. A. Scrosati at Baccaro Point.

We made the intertidal measurements described below at the high intertidal zone. For each location, we considered the intertidal range as the vertical distance between chart datum (0 m in elevation) and the highest elevation where perennial sessile organisms (the barnacle *Semibalanus balanoides*) occurred on the substrate outside of crevices (Scrosati and Heaven 2007). For each location, we divided the vertical intertidal range by three and collected the data described in the following paragraphs just above the bottom boundary of the upper third of the intertidal range. Since tidal amplitude increases from 1.8 m in L1 to 2.4 m in L9 (Tide-Forecast 2017), this method allowed us to take intertidal data along the coast at comparable elevations in terms of exposure to aerial conditions during low tides.

### Barnacle recruitment

For barnacles, recruitment refers to the appearance of new organisms in a benthic habitat after metamorphosis of settled pelagic larvae. On the Atlantic coast of Nova Scotia, *Semibalanus balanoides* is the only species of intertidal barnacle. It is a cross-fertilizing hermaphrodite that broods once per year (Bousfield 1954). In Atlantic Canada, this species mates in autumn, broods in winter, and releases pelagic larvae in spring (Bousfield 1954, Crisp 1968, Bouchard and Aiken 2012). On the studied coast, larvae settle in intertidal habitats in May and June (Ellrich et al. 2016a). As settled larvae quickly metamorphose into benthic recruits, the barnacle recruitment season spans May-June every year (Ellrich et al. 2015a). Thus, we measured barnacle recruitment at our nine locations in late June (Table 2).

**Table 2.**
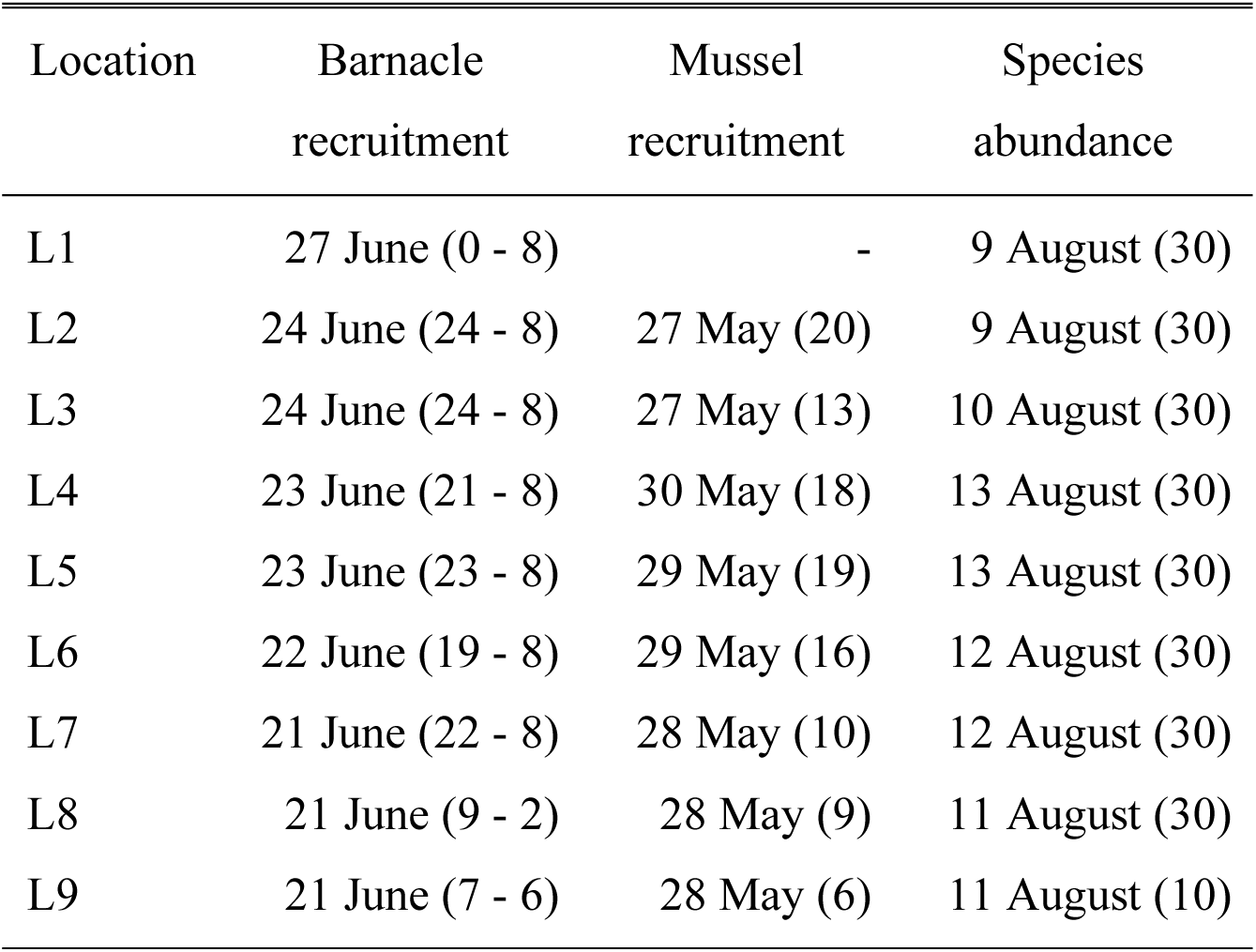
Dates when barnacle recruitment, mussel recruitment, and the abundance of barnacles, mussels, and dogwhelks were measured at the nine studied locations. Sample sizes are given in parentheses (for barnacle recruitment, the first number is the number of Safety-Walk tiles and the second number is the number of substrate clearings).

| Location | Barnacle<br>recruitment | Mussel<br>recruitment | Species<br>abundance |
| --- | --- | --- | --- |
| L1 | 27 June (0 - 8) | - | 9 August (30) |
| L2 | 24 June (24 - 8) | 27 May (20) | 9 August (30) |
| L3 | 24 June (24 - 8) | 27 May (13) | 10 August (30) |
| L4 | 23 June (21 - 8) | 30 May (18) | 13 August (30) |
| L5 | 23 June (23 - 8) | 29 May (19) | 13 August (30) |
| L6 | 22 June (19 - 8) | 29 May (16) | 12 August (30) |
| L7 | 21 June (22 - 8) | 28 May (10) | 12 August (30) |
| L8 | 21 June (9 - 2) | 28 May (9) | 11 August (30) |
| L9 | 21 June (7 - 6) | 28 May (6) | 11 August (10) |

To measure barnacle recruitment, we used two methods that provided bare substrate shortly before the recruitment season (in late April). One method used PVC tiles (8.9 cm × 4.6 cm × 0.4 cm) covered by grey Safety-Walk tape (3M, St. Paul, Minnesota, USA; pictured in Menge et al. 2010). This rubberized vinyl tape has a rugose texture and has been used to measure barnacle recruitment on other shores (Menge 2000, Lagos et al. 2008, Mazzuco et al. 2015). We attached each tile to the rocky substrate by screwing it to a hole previously drilled into the substrate. The other method involved clearing the natural rocky substrate (Bryson et al. 2014). We produced each clearing (10 cm × 10 cm) by removing all algae and invertebrates from the substrate using a chisel and a metallic mesh scourer. The number of tiles and clearings established at each location is specified in Table 2. We measured recruit density (recruitment) on the tiles and clearings by analyzing digital pictures of them taken in late June (Table 2).

### Mussel recruitment

Two mussel congeners occur in rocky intertidal habitats on the Atlantic coast of Nova Scotia, *Mytilus edulis* and *M. trossulus* (Tam and Scrosati 2011, 2014). These species show only subtle morphological differences (Innes and Bates 1999) and can even form hybrids (Riginos and Cunningham 2005), so their visual identification is difficult, especially at the recruit stage. Thus, we counted mussel recruits as *Mytilus* spp., as done in other field studies with these species (Cusson and Bourget 2005, Le Corre et al. 2013) and other mussel species (Wieters et al. 2008). To measure mussel recruitment, we installed plastic mesh scourers (Our Compliments Poly Pot Scrubbers, Mississauga, Ontario, Canada) at each location in late April and collected them at the end of May (Table 2) to measure recruit density in the lab. Mesh scourers have often been used to measure intertidal mussel recruitment (Menge and Menge 2013, Mazzuco et al. 2015, South 2016), as the scourers resemble habitats where the pelagic mussel larvae preferentially settle (filamentous algae or byssal threads of established mussels; Menge 1992; Le Corre et al. 2013). For *Mytilus edulis* and *M. trossulus*, pelagic pediveliger larvae of at least about 0.25 mm in shell length settle in those habitats and undergo metamorphosis, becoming benthic recruits (Bayne 1965, Menge et al. 2009, Martel et al. 2014). After growing to a shell length of about 0.5 mm (Hunt and Scheibling 1996, Le Corre et al. 2013), such recruits may enter a second pelagic dispersal phase (Bayne 1964). For instance, recruits of *M. edulis* up to 2.5 mm long can passively drift in the water aided by a byssus thread (Sigurdsson et al. 1976). Microscope observations indicated that approximately 70–80% of the recruits found in our scourers belonged to the first phase. Accurate estimates were not possible because the precise size threshold between both phases is not known (Le Corre et al. 2013). Nonetheless, since all of those organisms contribute to mussel recruitment (Le Corre et al. 2013), we counted the recruits of both phases together to determine recruit density for each scourer, as often done in field studies of this kind (Menge and Menge 2013).

### *Chlorophyll*-a *concentration (Chl*-a*) and particulate organic carbon (POC)*

We used MODIS-Aqua satellite data on Chl-*a* and POC for the 4 km × 4 km cells that include our nine locations (National Aeronautics and Space Administration 2017). Chl-*a* data are often used as a proxy for phytoplankton abundance (Menge and Menge 2013). Coastal Chl-*a* data from satellites are frequently used in intertidal ecology (Navarrete et al. 2005, Burrows et al. 2010, Arribas et al. 2014, Mazzuco et al. 2015, Lara et al. 2016). Small differences between satellite and in-situ data may exist. However, satellite data were the only available descriptors of Chl-*a* and POC for our locations. These data should be adequate for at least two reasons. On the one hand, there were clear differences in Chl-*a* and POC among locations (see Results), suggesting that the signal was stronger than potential noise. On the other hand, the data analyses identified Chl-*a* and POC as potentially important for barnacle and mussel recruitment and, on wave-exposed shores, pelagic larvae respond to Chl-*a* and POC conditions across extensive areas covered by the satellite cells. Overall, satellite data are practical when studying large extents of coastline (Legaard and Thomas 2006).

To evaluate potential Chl-*a* and POC forcing on barnacle and mussel recruitment, we used the monthly means of Chl-*a* and POC for April and May and (for barnacle recruitment) the June mean calculated from the beginning of that month and the dates when we measured barnacle recruitment (Table 2). For barnacles, we used May means of Chl-*a* and POC because recruits start to appear in early May (Ellrich et al. 2015a) and April means because the nauplius larvae of *S. balanoides* develop over 5–6 weeks feeding in coastal waters (Bousfield 1954, Drouin et al. 2002) before reaching the settling cyprid stage. A study done in 2013 at a nearby location revealed that most of the barnacle recruits appear during May. As dead recruits (indicated by empty shells on the substrate) were rare in June, most of the larvae that generated the June values of recruit density were likely in the water column in April (Scrosati and Ellrich 2016).

### Sea surface temperature (SST)

We measured SST using in-situ temperature loggers (HOBO Pendant Logger, Onset Computer, Bourne, Massachusetts, USA). At each location in late April (except at L1, where loggers were not deployed), we secured four loggers to the intertidal substrate using eye screws and cable ties. The loggers measured temperature every 30 min. We collected them on the dates when we measured barnacle recruitment (Table 2). As temperature was highly correlated between the loggers within locations (*r* = 0.84-0.99), for each location we generated a single time series of temperature by averaging the corresponding half-hourly values from the replicate loggers. Using these time series, we extracted for each location the values of daily SST for the study period, which we considered to be the temperature recorded at the time of the highest tide of each day. We determined the time of such tides using online information available for the closest locations to our studied locations (Table 1, Tide and Current Predictor 2017).

### Barnacle, mussel, and dogwhelk abundance

We measured the abundance of barnacles, mussels, and dogwhelks in natural communities in August (Table 2). At each location, we surveyed 30 random quadrats (20 cm × 20 cm), except at L9, where we only measured 10 quadrats. As often done in intertidal studies (Navarrete and Manzur 2008, Bryson et al. 2014), we quantified the abundance of barnacles and mussels as percent cover (using a 20 cm × 20 cm frame divided in 100 squares with monofilament line) and the abundance of dogwhelks as density (number of organisms per quadrat divided by quadrat area). We took these measurements during low tides.

### Data analyses

We evaluated whether the spring recruitment of barnacles and mussels and the summer abundance of barnacles, mussels, and dogwhelks differed among locations through separate analyses of variance (Sokal and Rohlf 2012). We searched for evidence of benthic-pelagic coupling following a model selection approach using the location averages of intertidal recruitment and pelagic traits. Specifically, we considered either barnacle recruitment or mussel recruitment as the dependent variable and the monthly means of Chl-*a*, POC, or SST as independent variables. Separately for these three pelagic traits, we compared all possible linear models (representing all possible combination of months) using their values of the corrected Akaike’s information criterion (AIC_c_). For each set of models, we considered the most plausible model as that exhibiting the lowest AIC_c_ score. Any other model within the set exhibiting a difference in AIC_c_ (∆AIC_c_) of less than 2 relative to the most plausible model was also considered to have substantial support (Burnham and Anderson 2004). The Results section also provides the relative likelihood of each model within any given set (Anderson 2008). We searched for evidence of bottom-up forcing by separately evaluating the relationships between dogwhelk abundance and the recruitment and abundance of barnacles and mussels (Sokal and Rohlf 2012). We conducted the analyses with JMP 9.0.

## Results

### Barnacle recruitment

Combining all locations, barnacle recruitment was lower on tiles covered by Safety-Walk tape than on cleared natural substrate (substrate type effects: *F*_1,189_ = 24.84, *P* < 0.001). This pattern was true especially at the locations with high recruitment (interaction term: *F*_7,189_ = 8.34, *P* < 0.001, Fig. 3). Therefore, we used the data from the substrate clearings to do the analyses on recruitment reported below. Based on such data, barnacle recruitment differed significantly along the Nova Scotia coast (*F*_8,55_ = 8.74, *P* < 0.001), peaking at one northern location (L2) and two southern locations (L7 and L8; Fig. 3).

**Figure 3.**
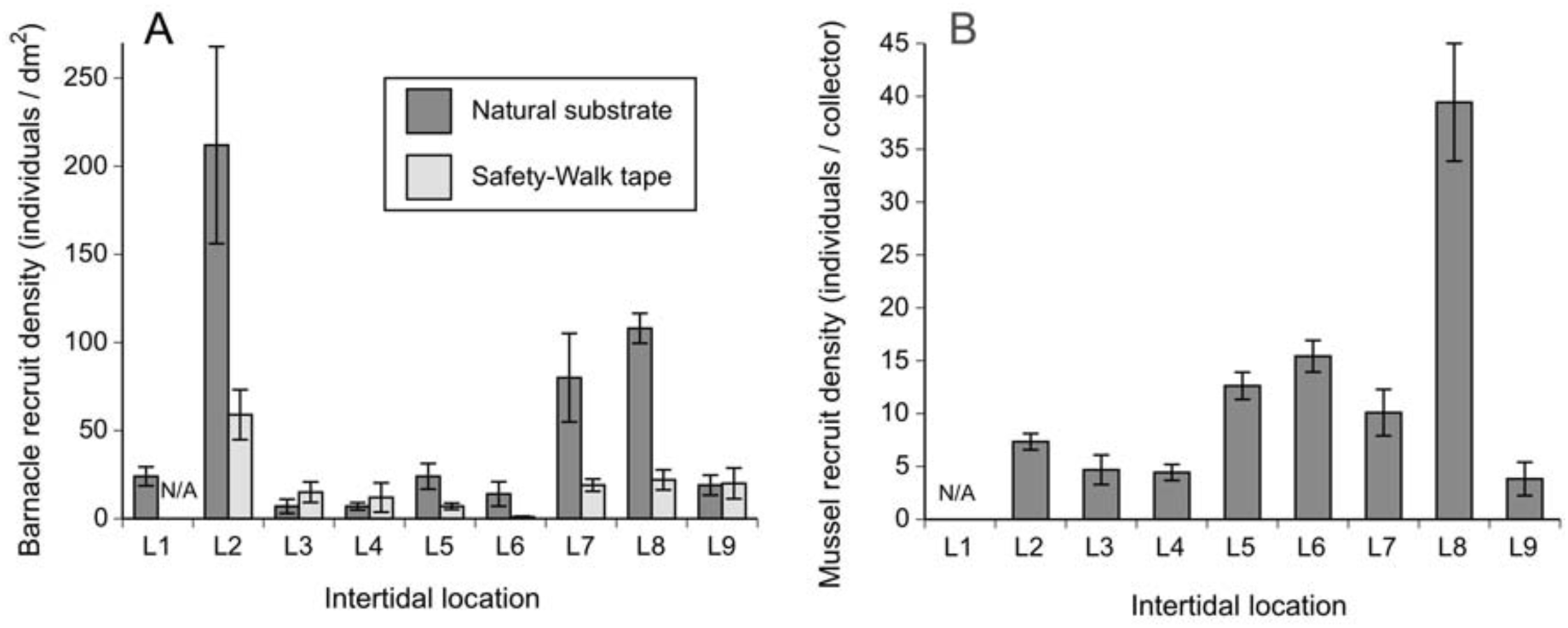
(A) Barnacle recruitment and (B) mussel recruitment (mean ± SE; see Table 2 for sample sizes) along the Atlantic coast of Nova Scotia.

Both Chl-*a* and POC also varied along the coast (Fig. 4). Model selection based on AIC_c_ scores revealed that the best model relating Chl-*a* with barnacle recruitment was the one including only April Chl-*a*, a positive relationship that explained 46% (adjusted *R*^2^ = 0.46) of the observed variation in recruitment (Table 3, Fig. 5). A similar result was obtained for POC, as the model including only April POC was the best one, also a positive relationship that explained 48% of the variation in recruitment (Table 3, Fig. 5). The AIC_c_ scores of the April Chl-*a* model and the April POC model were similar (Table 3), while April Chl-*a* and April POC were positively correlated, although not significantly (*r* = 0.54, *P* = 0.132).

**Figure 4.**
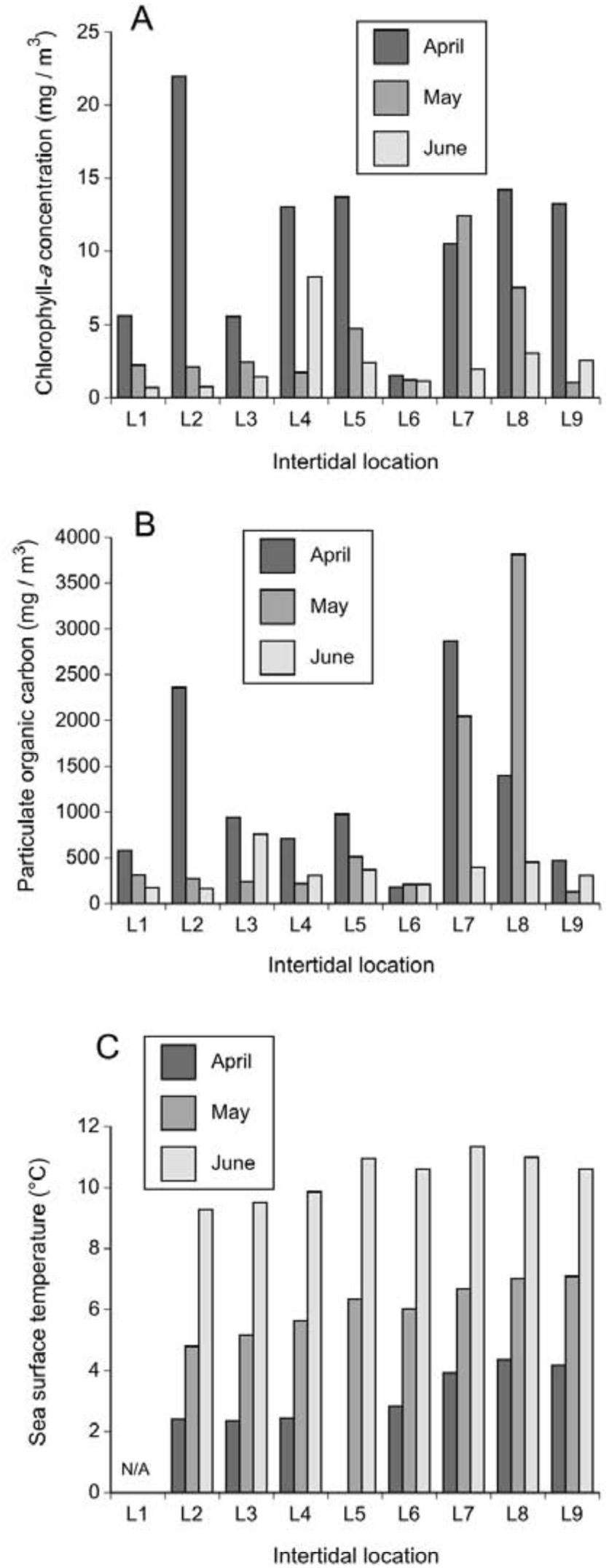
(A) Chlorophyll-*a* concentration, (B) particulate organic carbon, and (C) sea surface temperature along the Atlantic coast of Nova Scotia.

**Table 3.**
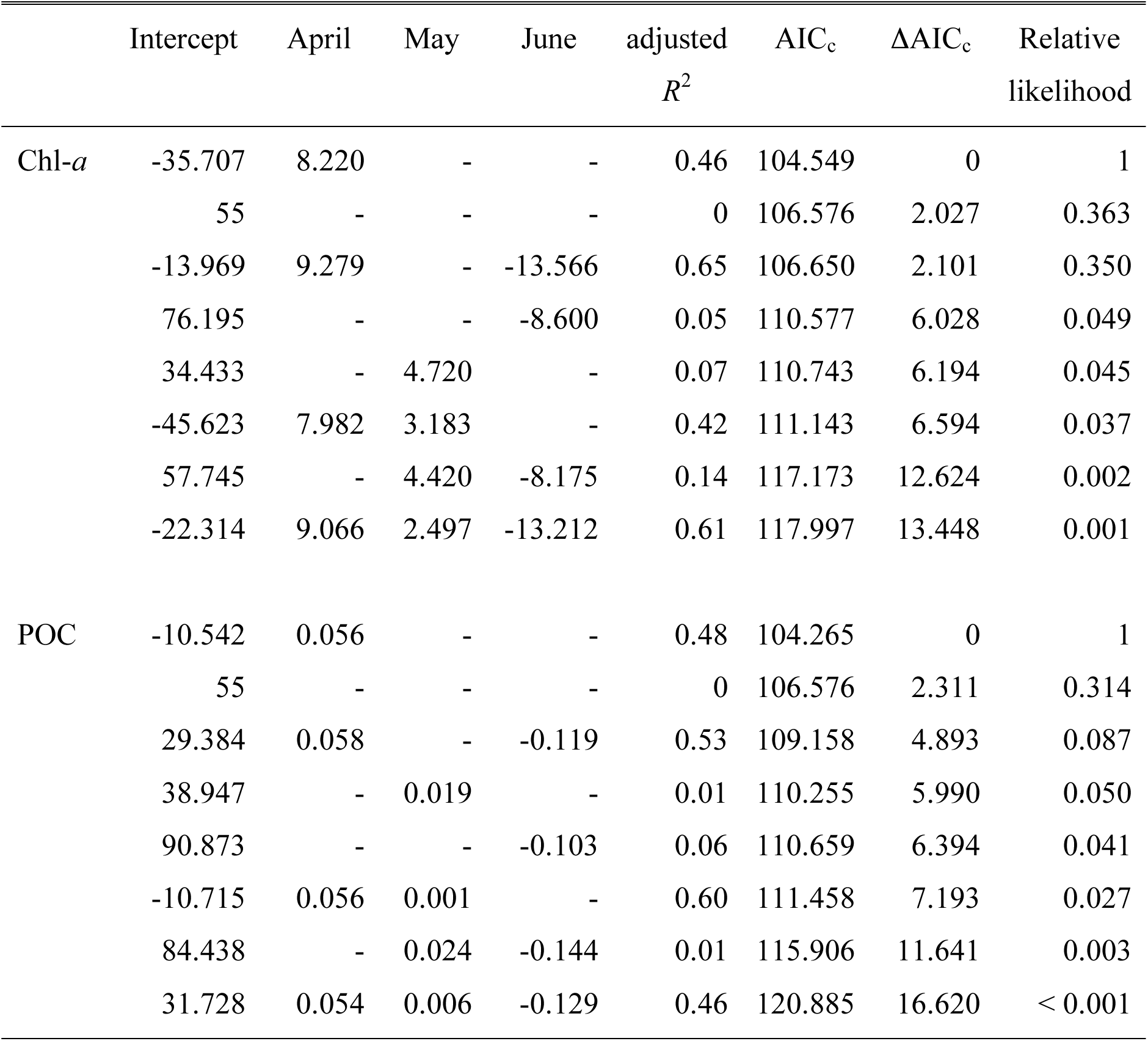
Summary information for the models relating barnacle recruitment and the April, May, and June means of Chl-*a* and POC. The numbers shown in the third, fourth, and fifth columns are the regression coefficients for April, May, and June, respectively.

| | Intercept | April | May | June | adjusted<br>$R^2$ | AIC<sub>c</sub> | ΔAIC<sub>c</sub> | Relative<br>likelihood |
| --- | --- | --- | --- | --- | --- | --- | --- | --- |
| Chl-*a* | -35.707 | 8.220 | - | - | 0.46 | 104.549 | 0 | 1 |
|  | 55 | - | - | - | 0 | 106.576 | 2.027 | 0.363 |
|  | -13.969 | 9.279 | - | -13.566 | 0.65 | 106.650 | 2.101 | 0.350 |
|  | 76.195 | - | - | -8.600 | 0.05 | 110.577 | 6.028 | 0.049 |
|  | 34.433 | - | 4.720 | - | 0.07 | 110.743 | 6.194 | 0.045 |
|  | -45.623 | 7.982 | 3.183 | - | 0.42 | 111.143 | 6.594 | 0.037 |
|  | 57.745 | - | 4.420 | -8.175 | 0.14 | 117.173 | 12.624 | 0.002 |
|  | -22.314 | 9.066 | 2.497 | -13.212 | 0.61 | 117.997 | 13.448 | 0.001 |
| POC | -10.542 | 0.056 | - | - | 0.48 | 104.265 | 0 | 1 |
|  | 55 | - | - | - | 0 | 106.576 | 2.311 | 0.314 |
|  | 29.384 | 0.058 | - | -0.119 | 0.53 | 109.158 | 4.893 | 0.087 |
|  | 38.947 | - | 0.019 | - | 0.01 | 110.255 | 5.990 | 0.050 |
|  | 90.873 | - | - | -0.103 | 0.06 | 110.659 | 6.394 | 0.041 |
|  | -10.715 | 0.056 | 0.001 | - | 0.60 | 111.458 | 7.193 | 0.027 |
|  | 84.438 | - | 0.024 | -0.144 | 0.01 | 115.906 | 11.641 | 0.003 |
|  | 31.728 | 0.054 | 0.006 | -0.129 | 0.46 | 120.885 | 16.620 | < 0.001 |

**Figure 5.**
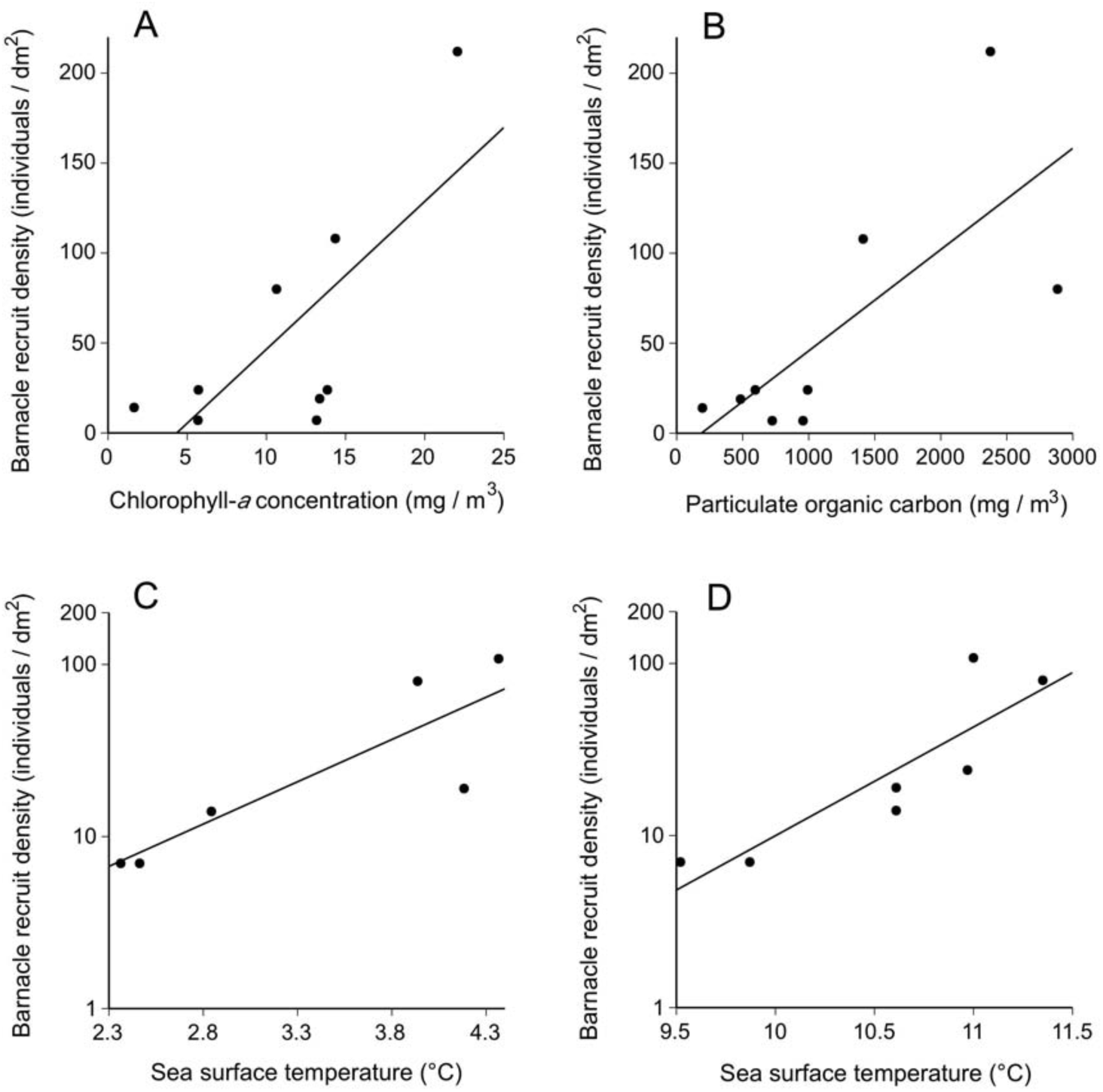
Relationships between barnacle recruitment and (A) April Chl-*a*, (B) April POC, (C) April SST, and (D) June SST, identified as relevant by AIC_c_-based model selection.

SST also varied along the coast (Fig. 4). AIC_c_-based model selection did not identify any model including SST as suitable to explain barnacle recruitment (Table 4). However, because of the seemingly overwhelming influence of Chl-*a* on recruitment in L2 (Figs 3–4), we explored excluding L2 from the SST analyses and applied a log_10_ transformation to the recruitment data. With this approach, model selection separately identified June SST and April SST as suitable explanatory variables through positive relationships (Fig. 5), although the model including only the intercept was similarly likely (Table 4).

**Table 4.** Summary information for the models relating barnacle recruitment and the April, May, and June means of SST and (excluding L2) log_10_ SST. The numbers shown in the third, fourth, and fifth columns are the regression coefficients for April, May, and June, respectively.

| | Intercept | April | May | June | adjusted<br>$R^2$ | AIC<sub>c</sub> | ΔAIC<sub>c</sub> | Relative<br>likelihood |
| --- | --- | --- | --- | --- | --- | --- | --- | --- |
| SST | 63.857 | - | - | - | 0 | 86.509 | 0 | 1 |
|  | 192.324 | - | -21.204 |  | 0.13 | 93.054 | 6.545 | 0.038 |
|  | 55.917 | 2.470 | - | - | 0.20 | 93.503 | 6.994 | 0.030 |
|  | 242.667 | - | - | -17.341 | 0.16 | 93.290 | 6.781 | 0.034 |
|  | 765.093 | 229.960 | -237.772 | - | 0.68 | 96.801 | 10.292 | 0.006 |
|  | 613.914 | 53.666 |  | -70.077 | 0.29 | 106.432 | 19.923 | < 0.001 |
|  | 51.613 |  | -40.251 | 24.837 | 0.39 | 106.961 | 20.452 | < 0.001 |
|  | 598.801 | 230.897 | -261.480 | 29.764 | 0.60 | 138.200 | 51.691 | < 0.001 |
| $\log_{10}$ SST | -5.758 | - | - | 0.677 | 0.77 | 15.944 | 0 | 1 |
|  | 1.342 | - | - | - | 0 | 15.984 | 0.040 | 0.980 |
|  | -0.304 | 0.492 | - | - | 0.68 | 17.824 | 1.880 | 0.391 |
|  | -1.931 | - | 0.522 | - | 0.54 | 20.044 | 4.100 | 0.129 |
|  | -1.677 | 0.946 | -0.952 | 0.555 | 0.88 | 39.894 | 23.950 | < 0.001 |
|  | -4.058 | 0.21 | - | 0.448 | 0.76 | 44.404 | 28.460 | < 0.001 |
|  | -5.451 |  | 0.082 | 0.599 | 0.70 | 45.797 | 29.853 | < 0.001 |
|  | 1.919 | 1.037 | -0.646 | - | 0.65 | 46.606 | 30.662 | < 0.001 |

### Mussel recruitment

Mussel recruitment varied significantly along the coast (F_7,103_ = 30.55, *P* < 0.001), exhibiting a peak at a southern location where barnacle recruitment also peaked (L8; Fig. 3). Model selection based on AIC_c_ scores did not identify any model including Chl-*a* data as suitable to explain mussel recruitment (Table 5). However, using POC data, model selection indicated that the best model included May POC, a positive relationship that explained 70% of the observed variation in mussel recruitment (Table 5, Fig. 6). None of the models using SST data was suitable to explain mussel recruitment (Table 5) and no data transformations such as those applied to barnacle recruitment modified this outcome.

**Table 5.** Summary information for the models relating mussel recruitment and the April and May means of Chl-*a*, POC, and SST. The numbers shown in the third and fourth columns are the regression coefficients for April and May, respectively.

| | Intercept | April | May | adjusted<br>$R^2$ | AIC<sub>c</sub> | ΔAIC<sub>c</sub> | Relative<br>likelihood |
| --- | --- | --- | --- | --- | --- | --- | --- |
| Chl-*a* | 12.239 | - | - | 0 | 67.463 | 0 | 1 |
|  | 7.281 | - | 1.195 | 0.03 | 71.624 | 4.161 | 0.125 |
|  | 11.687 | 0.582 | - | < 0.01 | 73.058 | 5.595 | 0.061 |
|  | 7.347 | -0.006 | 1.196 | 0.17 | 80.957 | 13.494 | 0.001 |
| POC | 5.12 | - | 0.008 | 0.70 | 62.083 | 0 | 1 |
|  | 9.525 | -0.005 | 0.009 | 0.80 | 66.945 | 4.862 | 0.088 |
|  | 12.239 | - | - | 0 | 67.463 | 5.380 | 0.068 |
|  | 11.284 | 0.001 | - | 0.16 | 73.033 | 10.950 | 0.004 |
| SST | 12.183 | - | - | 0 | 61.364 | 0 | 1 |
|  | -12.153 | 7.569 | - | 0.14 | 66.001 | 4.637 | 0.098 |
|  | -27.498 | - | 6.550 | 0.06 | 66.643 | 5.279 | 0.071 |
|  | 5.298 | 13.167 | -5.85 | 0.05 | 79.830 | 18.466 | < 0.001 |

**Figure 6.**
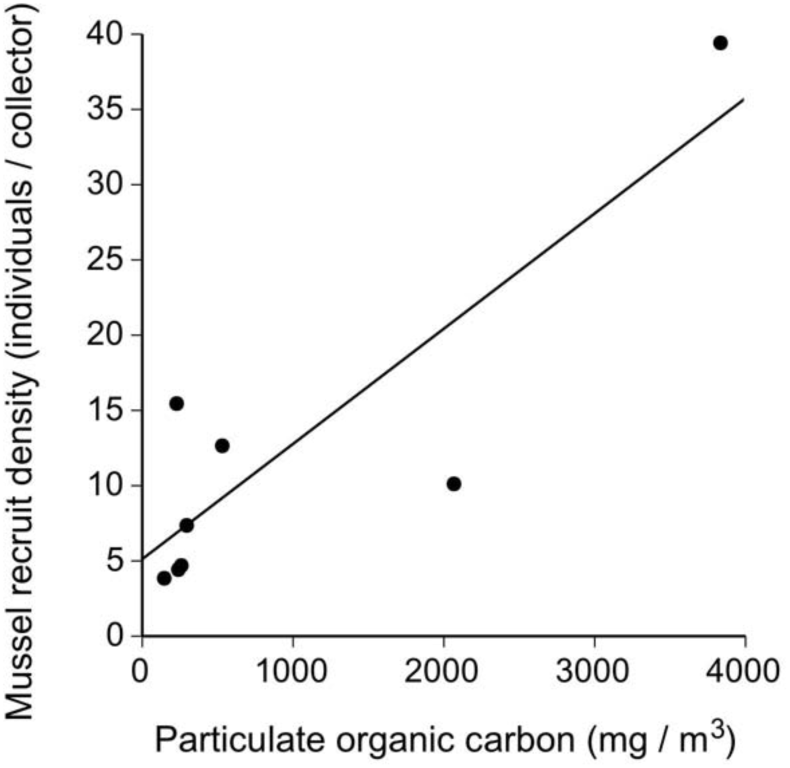
Relationship between mussel recruitment and May POC, identified as relevant by AIC_c_-based model selection.

### Barnacle and mussel abundance

Barnacle abundance measured in pristine communities in August varied significantly along the coast (*F*_8, 241_ = 38.58, *P* < 0.001; Fig. 7). The August abundance of barnacles was positively related to barnacle recruitment measured in June (adjusted *R*^2^ = 0.82, *P* < 0.001; Fig. 7). Mussel abundance measured in pristine communities in August also varied significantly along the coast (F_8, 241_ = 55.51, *P* < 0.001; Fig. 7). The August abundance of mussels was positively related to mussel recruitment measured in May (adjusted *R*^2^ = 0.87, *P* < 0.001; Fig. 7).

**Figure 7.**
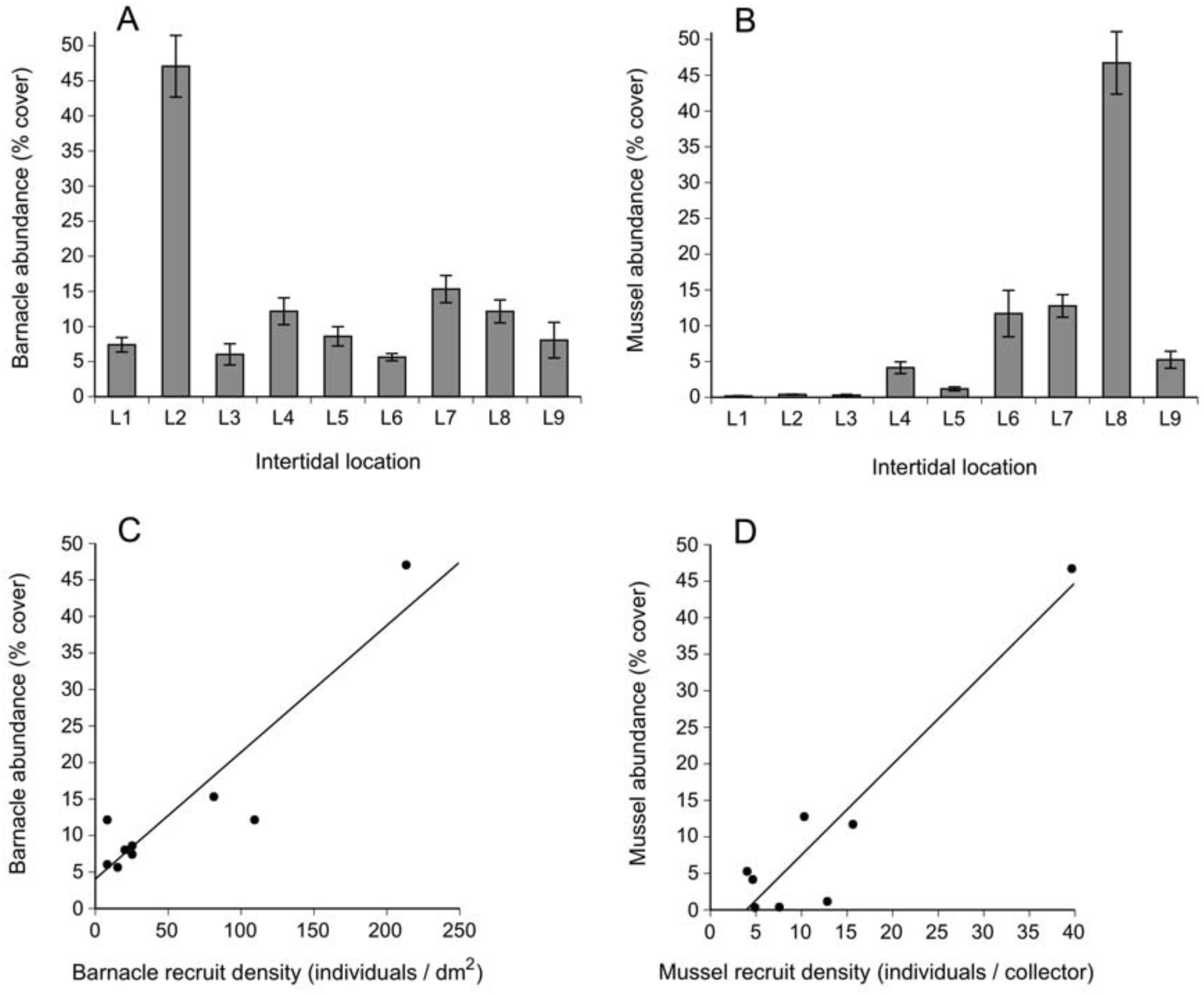
Summer abundance of (A) barnacles and (B) mussels (mean ± SE; see Table 2 for sample sizes) along the Atlantic coast of Nova Scotia and relationships with the spring recruitment of (C) barnacles and (D) mussels.

### Dogwhelk abundance

Dogwhelk abundance measured in pristine communities in August varied significantly along the coast (*F*_8, 241_ = 6.48, *P*, 0.001; Fig. 8). Dogwhelk abundance was positively related to barnacle recruitment (adjusted *R*^2^ = 0.70, *P* = 0.003) and barnacle abundance (adjusted *R*^2^ = 0.45, *P* = 0.029; Fig. 8), but unrelated to mussel recruitment (adjusted *R*^2^ = 0.20, *P* = 0.150) or mussel abundance (adjusted *R*^2^ = 0.09, *P* = 0.222). However, because of intense ice scour at the four northernmost locations (L1 to L4) in early April (Petzold et al. 2014), the intertidal substrate at those locations did not have any organisms (filamentous algae or mature mussels) capable of attracting mussel recruits during the period when we measured recruitment using mesh collectors. Thus, the obtained values of mussel recruitment for those four locations are best seen as potential recruitment. After excluding the mussel recruitment data for those four locations, the positive correlation between dogwhelk abundance and mussel recruitment became significant (adjusted *R*^2^ = 0.93, *P* = 0.005; Fig. 8), a remarkable outcome given that this result was based merely on the remaining five locations (L5 to L9).

**Figure 8.**
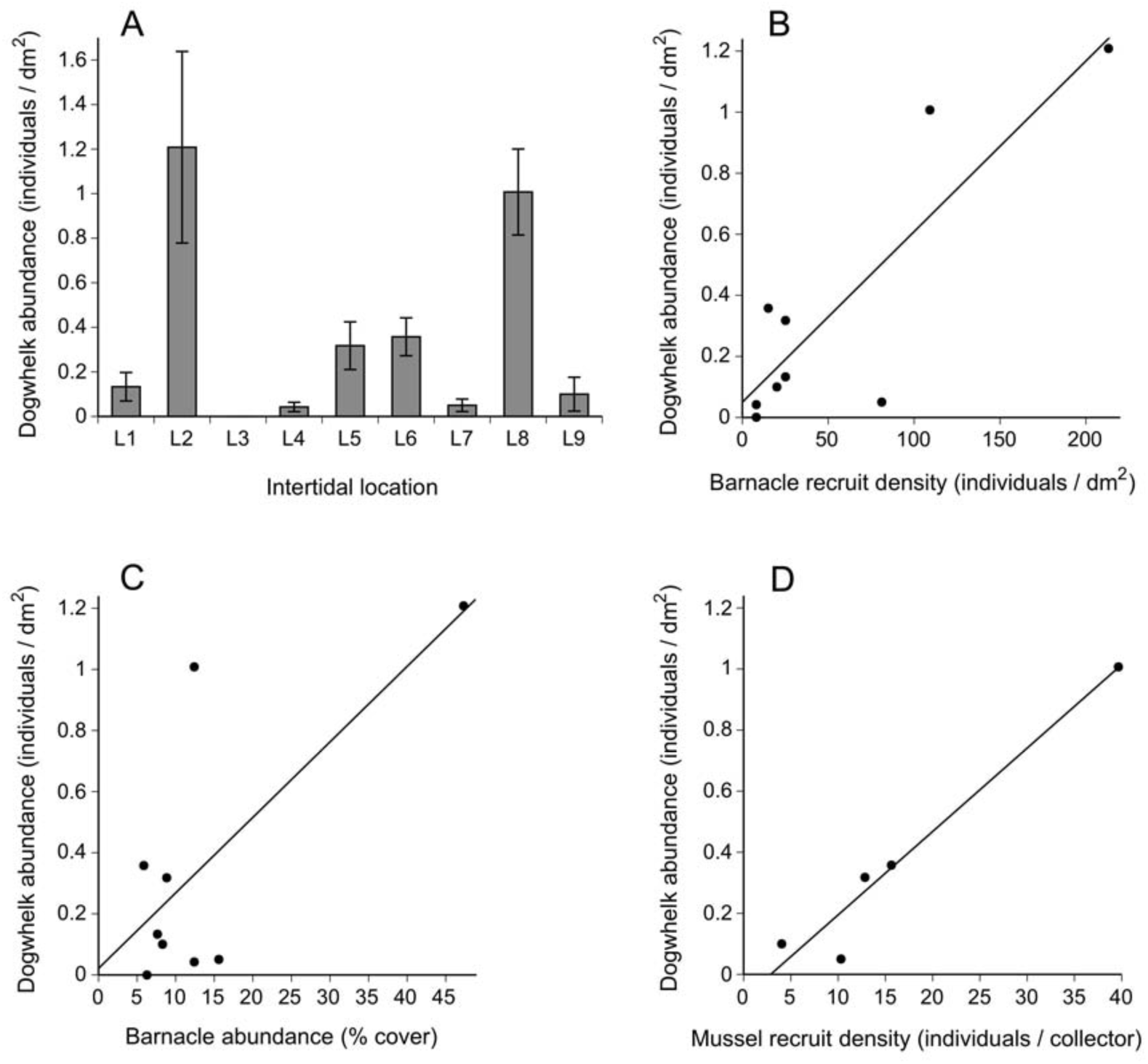
(A) Dogwhelk summer abundance (mean ± SE; see Table 2 for sample sizes) along the Atlantic coast of Nova Scotia and relationships with (B) barnacle recruitment, (C) barnacle abundance, and (D) mussel recruitment.

## Discussion

### Benthic-pelagic coupling

Our study has revealed that intertidal barnacle and mussel recruitment varied markedly along the Atlantic coast of mainland Nova Scotia. Differences were not correlated to latitude, however, as recruitment peaked in southern and northern locations. The data analyses indicated that pelagic food supply was likely influential for intertidal recruitment, as it would be expected given that barnacles and mussels are filter-feeders.

For barnacles in our study, both phytoplankton abundance (Chl-*a*) and particulate organic carbon were identified as relevant by the model selection approach. It is not possible to tell, however, if both pelagic traits were biologically relevant or simply collinear. Phytoplankton is the main food source for barnacle nauplius larvae (Walker et al. 1987, Turner et al. 2001), so the observed relationship with Chl-*a* probably reflects a true biological effect on recruitment through influences on larval nutrition. Another study done in Nova Scotia concluded that most of the barnacle larvae that make up recruit counts in June are in the water column in April (Scrosati and Ellrich 2016). This could explain why the best model relating Chl-*a* with barnacle recruitment only included April Chl-*a* data. Interestingly, Barnes (1956) found that the development of *S. balanoides* larvae on a UK shore improved with planktonic diatom abundance. For mussels, only POC (not Chl-*a*) was identified by our data analyses as potentially relevant for recruitment. The best model included only May POC, suggesting that particulate organic carbon might be important for mussel recruitment by influencing late-stage larvae and perhaps early recruits. Support for this notion exists for freshwater mussels (Nalepa et al. 1995).

Given these findings, an emerging question of interest is why pelagic food supply varied along the coast. The intermittent upwelling hypothesis (IUH; Menge and Menge 2013) posits that nearshore phytoplankton abundance can be influenced by coastal upwelling. Frequent, intense upwelling would drive upwelled inorganic nutrients offshore, preventing the occurrence of phytoplankton blooms near the coast. Intermittent upwelling would allow for upwelled nutrients to stay near the shore for longer periods, fueling a higher phytoplankton abundance (Menge and Menge 2013). Upwelling has been reported for Nova Scotia (Tee et al. 1993), but that study only covered offshore areas in the south. Thus, it is not currently possible to ascertain if possible upwelling differences along the Nova Scotia coast (yet to be determined) could drive latitudinal changes in pelagic food supply. The IUH also predicts that intermittent upwelling increases intertidal filter-feeder recruitment also by enhancing the coastal retention of pelagic larvae (Menge and Menge 2013). This upwelling-recruitment relationship has been supported by data from Oregon, California, and New Zealand (Menge and Menge 2013). Recently, however, alternative approaches failed to support the IUH and suggested that surf zone width and tidally generated internal waves are more important for coastal phytoplankton abundance and intertidal recruitment (Shanks et al. 2017a,b, Shanks and Morgan 2017). Clearly, the oceanographic factors driving pelagic food supply and filter-feeder recruitment are currently under debate. Thus, those studies suggest potentially useful approaches to understand the oceanographic causes of the benthic-pelagic coupling observed on the Nova Scotia coast.

Sea surface temperature also varied along the Nova Scotia coast, but its relevance to intertidal recruitment was less clear. Seawater temperature can limit *S. balanoides* recruitment, although under relatively high temperature ranges, as seen on European shores (Abernot-Le Gac et al. 2013, Rognstad et al. 2014). On the Gulf of St. Lawrence coast, which exhibits low temperatures in spring (this gulf freezes in winter), seawater temperature was positively related to *S. balanoides* recruitment, presumably through the enhancement of larval performance (Scrosati and Ellrich 2016). Thus, it was not surprising to find positive relationships between *S. balanoides* recruitment and SST in our study. However, the model selection approach also indicated that no SST-recruitment relationship was similarly likely. Overall, these results suggest that pelagic food supply might have a larger influence on barnacle recruitment than seawater temperature on the Atlantic coast of Nova Scotia. This notion is further supported by the larger variation (coefficient of variation based on location means) that April Chl-*a* and POC exhibited among locations (55% and 77%, respectively) than either April, May, or June SST (28%, 14%, and 7%, respectively). For mussels, the model selection approach did not detect any plausible SST–recruitment relationship, leaving POC as the only studied pelagic trait with a possible influence, further supporting the seemingly relevant role of pelagic food supply for intertidal filter-feeder recruitment on this coast.

On the NE Atlantic coast, *S. balanoides* recruit density can be similar or higher (Hawkins and Hartnoll 1982, Kendall et al. 1985, Jenkins et al. 2000, 2008, Kent et al. 2003, Rognstad et al. 2014) than reported here for the Nova Scotia coast. The fact that, in combination, those studies surveyed a wider range of wave exposure, intertidal elevation, and food supply than our study may explain the higher recruitment range that they reported. On the NE Pacific coast, intertidal barnacle recruitment is often high. For example, in Oregon and California, recruits of *Balanus glandula* and *Chthamalus dalli* appear throughout most of the year and can reach, when combined, mean densities of 1800 recruits/dm^2^ in one month (Navarrete et al. 2008). On that coast, either intermittent upwelling or surf zone properties would allow for barnacle larvae to remain near the coast and would favor high coastal Chl-*a* (above 23 mg/m^3^), enhancing intertidal barnacle recruitment (Menge and Menge 2013, Shanks and Morgan 2017). The low SST found on the Nova Scotia coast in April might further limit recruit density relative to the NE Pacific coast. The higher recruitment reported for the NE Atlantic (Jenkins et al. 2000) and NE Pacific (Navarrete et al. 2008) coasts might also result from the elevation where data were taken. We surveyed the bottom of the upper third of the vertical intertidal range, but those studies surveyed middle (Jenkins et al. 2000) and middle-to-low (Navarrete et al. 2008) elevations. On the Gulf of St. Lawrence coast of Nova Scotia, recruit density in 2006 was three times higher at middle elevations and two times higher at low elevations than at high elevations (MacPherson and Scrosati 2008). Thus, recruitment differences between these coasts, although present, could be smaller if data were available for the same relative elevation.

A comment on the methods to measure barnacle recruitment is pertinent. The tiles covered by Safety-Walk tape often yielded lower recruitment than cleared rocky substrates. A poor relative performance of Safety-Walk tape has also been reported for the Pacific coast (Shanks 2009). Nonetheless, such tiles have been used extensively on the Pacific coast of North and South America (Lagos et al. 2008, Menge et al. 2010). In high-recruitment Pacific shores, such tiles are quickly covered by barnacle recruits (R. A. Scrosati, pers. obs.), so substrate quality (tape vs. rock) might be unimportant for larval settlement choices, rendering Safety-Walk tape as an acceptable surface to measure recruitment. On the Atlantic coast of Nova Scotia, however, where recruitment is generally lower, Safety-Walk tape seems inadequate for that purpose. We note that recent field experiments that measured barnacle recruitment in Nova Scotia (Ellrich et al. 2015a,b, 2016a,b, Ellrich and Scrosati 2016) also used tiles covered by a rugose tape, but that tape (Permastik self-adhesive anti-skid safety tread, RCR International, Boucherville, Quebec, Canada) has a sandpaper texture and yielded similar recruitment values as the adjacent rock.

### Bottom-up forcing

On the Atlantic coast of Nova Scotia, dogwhelks are the main predators of intertidal barnacles and mussels (Hunt and Scheibling 1998, Ellrich et al. 2015a, Sherker et al. 2017). This seems to be especially true at the high-intertidal, wave-exposed habitats surveyed for this study. Sea stars (Keppel et al. 2015) and crabs (Boudreau et al. 2017) are also predators of mussels in Atlantic Canada. However, we never found such organisms in the surveyed habitats. At low intertidal elevations, where emersion-related abiotic stress is lower (Eckersley and Scrosati 2012), sea stars and crabs were sometimes present, but in very low abundances. It is only in wave-sheltered intertidal habitats where crabs are more common in Nova Scotia (Boudreau et al. 2017), although sea stars are still rare in such places (Scrosati and Heaven 2007), unlike on other shores of the world (Hayne and Palmer 2013). For these reasons, it was not surprising to find that dogwhelk abundance was correlated to barnacle recruitment and abundance and mussel recruitment in our study. It is remarkable that these relationships were based on a relatively limited number of locations, indicating the strength of these patterns.

Although our data are mensurative, it is worth speculating on possible underlying processes to orient future research. Our results suggest that barnacle recruitment is relevant for dogwhelks along the coast. This is especially apparent when examining northern locations. In early April (before we initiated field work), sea ice formed during the winter on the Gulf of St. Lawrence drifted out of that gulf and reached our four northernmost Atlantic locations (L1 to L4), causing extensive intertidal disturbance (Petzold et al. 2014). After ice melt (which occurred before late April), the main organisms that recolonized such locations were barnacles. This was most clear at L2, the location where barnacle recruits were most abundant and, by far, the main organisms during early post-ice succession. Only the seaweed *Fucus vesiculosus* recolonized that location relatively abundantly, but only later in the summer. Dogwhelks were not affected by ice scour, because these organisms stay in deep crevices during the cold season (Crothers 1985, Gosselin and Bourget 1989, R. A. Scrosati, pers. obs.) and only become active in the spring (Hughes 1972, Hunt and Scheibling 1998). Although many barnacle recruits reached adult sizes by the fall, many others were eaten by dogwhelks. Now, dogwhelk density in August was certainly not determined exclusively by barnacle recruitment in the previous spring, as most dogwhelks were older (based on their size). However, the growing barnacle recruits were the only food source for dogwhelks in the northern locations in 2014 because of the previous ice scour. Also, observations made just before ice scour indicated that mussels were then present, but rare, at such locations (Petzold et al. 2014). In addition, observations done in previous years for other purposes indicated that barnacle recruitment was always higher at L2 than at the other northern locations, resembling the pattern for 2014. In combination, these lines of evidence suggest an important role of barnacle recruitment for dogwhelk abundance. The field manipulation of barnacle recruit density could confirm this notion.

The correlation between barnacle spring recruitment and summer abundance suggests the persistence of the pelagic signal to some extent. A positive relationship between barnacle recruitment and adult abundance was also observed on the Pacific coast (Blanchette et al. 2006). On the other hand, the relationship between the summer abundance of barnacles and dogwhelks may also have resulted from barnacles being relevant for dogwhelks. However, the fact that abundance was measured at the same time (August) for both species suggests some caution for this inference.

Our results also suggest that mussel recruitment may also have been important for dogwhelks, although primarily at central and southern locations. At the northern locations, the biological landscape while we measured mussel recruitment was simple (mostly only barnacle recruits) because of the previous ice scour. Thus, for those locations, our mussel recruitment data (measured with collectors resembling complex habitats) indicate potential recruitment. Considering only the central and southern locations, mussel recruitment and dogwhelk abundance were strongly correlated, suggesting a relevance that field experiments could confirm. As the summer abundance of mussels was uncorrelated to that of dogwhelks, it could be thought that mussels did not influence dogwhelk abundance. However, under the view that barnacles were likely the main food source for dogwhelks at the northern locations (see above), excluding L1 to L4 yielded a significant correlation between mussel and dogwhelk abundance (adjusted *R*^2^ = 0.70, *P* = 0.049), suggesting a role for mature mussels at central and southern locations. As both such variables were measured at the same time, experimental confirmation is needed. We note that a larger study spanning 1800 km between Newfoundland and Long Island (although with a poorer coverage of Nova Scotia than the present study) also found a positive correlation between mussel and dogwhelk abundance (Tam and Scrosati 2011).

It is also worth discussing the correlation between mussel spring recruitment and summer abundance. Although summer abundance did not result exclusively from the growth of spring recruits (because many mussels were older, as per shell growth rings; Tam and Scrosati 2011), the correlation indicates the potential that, as noted for barnacles, the pelagic signal persists for some time during mussel growth. For other coasts of the world, mussel recruitment and abundance may (Blanchette et al. 2006, Arribas et al. 2014) or may not (Menge et al. 2004, Wieters et al. 2008) be correlated across locations.

The importance of barnacle and mussel recruitment for dogwhelks is further supported by *Nucella lapillus* being a direct developer without pelagic stages (Crothers 1985). Such a trait is expected to make dogwhelk abundance more dependent on local food sources than if pelagic dispersal were possible (Navarrete and Manzur 2008). In fact, dogwhelk abundance has also been found to correlate to barnacle and mussel recruitment on the Pacific coast (Wieters et al. 2008) and to mussel recruitment in New England (Menge 1976). Although sea stars do have pelagic larvae and thus higher dispersal than dogwhelks, bottom-up effects of mussel recruitment on sea star recruitment have been reported for subtidal habitats in Maine (Witman et al. 2003). Recent studies on rocky intertidal locations from the Gulf of Maine have further concluded that oceanographically driven variation in intertidal recruitment might explain the observed changes in community organization along that coast (Bryson et al. 2014).

Mensurative studies are valuable to investigate relatively unknown large ecosystems (Hughes et al. 2002, Sagarin and Pauchard 2010), as the Atlantic Canadian coast is in terms of benthic-pelagic coupling and bottom-up forcing. Overall, this study has revealed a geographic structure in filter-feeder recruitment and suggests that benthic-pelagic coupling and bottom-up forcing may ultimately influence intertidal communities. Thus, this contribution adds to the few equivalent studies done on western ocean boundary coasts (Menge and Menge 2013, Arribas et al. 2014, Mazzuco et al. 2015). We suggest that investigating oceanographic influences on pelagic food supply and experimentally evaluating the above discussed interspecific interactions should be relevant next steps along this research line.

## Acknowledgements.

This project was funded by grants awarded to Ricardo A. Scrosati by the Canada Research Chairs program, the Natural Sciences and Engineering Research Council (NSERC Discovery Grant), and the Canada Foundation for Innovation (Leaders Opportunity Grant) and by a postdoctoral scholarship awarded to Julius A. Ellrich by the German Academic Exchange Service (DAAD). We are grateful to Willy Petzold and Melanie Spieker for field and laboratory assistance.

